# Tau Protein Modulates an Epigenetic Mechanism of Cellular Senescence

**DOI:** 10.1101/2023.06.05.543662

**Authors:** Claudia Magrin, Martina Sola, Ester Piovesana, Marco Bolis, Andrea Rinaldi, Stéphanie Papin, Paolo Paganetti

**Author notes:** **Correspondence:** Prof. Paolo Paganetti, Laboratory for Aging Disorders, LRT EOC, Via Chiesa 5, 6500 Bellinzona, Switzerland. These authors share last authorship (Stéphanie Papin, Paolo Paganetti).

## Abstract

Progressive Tau deposition in neurofibrillary tangles and neuropil threads is the hallmark of tauopathies, a disorder group that includes Alzheimer’s disease. Since Tau is a microtubule-associated protein, a prevalent concept to explain the pathogenesis of tauopathies is that abnormal Tau modification contributes to dissociation from microtubules, assembly into multimeric β-sheets, proteotoxicity, neuronal dysfunction and cell loss. Tau also localizes in the cell nucleus and evidence supports an emerging function of Tau in DNA stability and epigenetic modulation. To better characterize the possible role of Tau in regulation of chromatin compaction and subsequent gene expression, we performed a bioinformatics analysis of transcriptome data obtained from Tau-depleted human neuroblastoma cells. Among the transcripts deregulated in a Tau-dependent manner, we found an enrichment of target genes for the polycomb repressive complex 2. We further describe decreased cellular amounts of the core components of the polycomb repressive complex 2 complex and a lower histone 3 trimethylation activity in Tau deficient cells. Among the de-repressed polycomb repressive complex 2 target gene products, IGFBP3 protein was found to be linked to increased senescence induction in Tau-deficient cells. Our findings propose a mechanism for Tau-dependent epigenetic modulation of cell senescence, a key event in pathologic aging.

## 1 Introduction

Tau pathology is the hallmark of tauopathies, a neurodegenerative disorder group that includes Alzheimer’s disease (AD), where progressive Tau deposition in neurofibrillary tangles and neuropil threads correlates with a deteriorating clinical course (Long and Holtzman, 2019). Autosomal dominant mutations in the *MAPT* gene encoding for Tau lead to a relatively small group of frontotemporal lobar degenerations (FTLD-Tau), which are classified among frontotemporal dementia (Josephs, 2018) diagnosed mostly at 45-65 years of age (Hutton et al., 1998, Spillantini et al., 1998). With Tau being a microtubule-binding protein, a prevalent concept to explain the pathogenesis of tauopathies is that abnormal Tau modification e.g., phosphorylation and folding, contributes to Tau dissociation from microtubules, assembly into multimeric β-sheets, proteotoxicity, neuronal dysfunction and cell loss (Jeganathan et al., 2006, Ludolph et al., 2009). In addition to its well-characterized role in neurodegeneration, studies reporting a correlation between *MAPT* gene products and survival in various types of tumors endorse an implication of Tau in cancer (Papin and Paganetti, 2020, Gargini et al., 2020, Gargini et al., 2019, Cimini et al., 2022, Rossi et al., 2018). The mechanisms underlying these findings may involve microtubule-unrelated Tau functions.

Tau exerts non canonical functions e.g., it localizes in the cell nucleus and binds DNA (Cross et al., 1996, Greenwood and Johnson, 1995, Loomis et al., 1990, Thurston et al., 1996). Heat or oxidative stress cause nuclear translocation of Tau, which may favor its role in DNA protection (Sultan et al., 2011). Neurons knocked-out for *MAPT* display enhanced DNA damage (Violet et al., 2014), and induced DNA damage correlates with nuclear translocation and dephosphorylation of Tau (Ulrich et al., 2018). Chromosomal abnormalities in AD fibroblasts (Rossi et al., 2008) and frequent DNA damage in AD brains (Lovell and Markesbery, 2007, Mullaart et al., 1990), both reinforce the emerging function of Tau in DNA stability. Tau depletion also modulates the induction of apoptosis and cell senescence in response to DNA damage by a mechanism involving P53, the guardian of the genome (Sola et al., 2020). Additional functions of Tau in epigenetic modulation were also reported (Rico et al., 2021). Upon binding to histones, Tau stabilizes chromatin compaction (Montalbano et al., 2021, Frost et al., 2014, Rico et al., 2021, Montalbano et al., 2020) and affects global gene expression during the neurodegenerative process (Frost et al., 2014, Klein et al., 2019). A meta-analysis of dysregulated DNA methylation in AD identified over hundreds genomic sites in cortical regions (Shireby et al., 2022); growing to thousands when looking at the dentate gyrus of oldest old patients (Lang et al., 2022).

DNA and histone modification is an effective mechanism to regulate gene activity. Hence, with the aim to investigate a possible participation of Tau in gene expression, we performed a bioinformatics analysis of transcriptome data obtained from Tau-depleted human neuroblastoma cells. Among the transcripts deregulated in a Tau-dependent manner, we found an enrichment of target genes for the Polycomb Repressive Complex 2 (PRC2), a result confirmed by decreased PRC2 protein and histone 3 (H3) methylation in Tau deficient cells. Notably, among the de-repressed gene products, Insulin Growth Factor Binding Protein 3 (IGFBP3) was linked to senescence induction. Our findings propose a Tau-driven mechanism for epigenetic modulation of cell senescence, a key event in pathologic aging.

## 2 Materials and methods

### 2.1 Cell culture

Human SH-SY5Y cells (94030304, Sigma-Aldrich) were cultured in complete Dulbecco’s Modified Eagle Medium (61965–059, Gibco) supplemented with 1% non-essential amino acids (11140035, Gibco), 1% penicillin-streptomycin (15140122, Gibco) and 10% fetal bovine serum (FBS; 10270106, Gibco). Cells were grown at 37°C with saturated humidity and 5% CO_2_ and maintained in culture for less than one month. *MAPT* knock-out cells were as described (Sola et al., 2020).

### 2.2 RNAseq

Total RNA extraction with the TRIzol™ Reagent (15596026, Invitrogen) was done according to the instructions of the manufacturer. Extracted RNA was processed with the NEBNext Ultra Directional II RNA library preparation kit for Illumina and sequenced on the Illumina NextSeq500 with single-end, 75 base pair long reads. The overall quality of sequencing reads was evaluated using a variety of tools, namely FastQC (Wingett and Andrews, 2018), RSeQC (Wang et al., 2012), AfterQC (Chen et al., 2017) and Qualimap (García-Alcalde et al., 2012). Sequence alignments to the reference human genome (GRCh38) was performed using STAR (v.2.5.2a) (Dobin et al., 2013). Transcript expression was quantified at gene level with the comprehensive annotations v27 release of the Gene Transfer File (GTF) made available by Gencode (Harrow et al., 2012). Raw-counts were further processed in the R Statistical environment and downstream differential expression analysis was performed using the DESeq2 pipeline (Love et al., 2014). Transcripts characterized by low mean normalized counts were filtered out by the Independent Filtering feature embedded in DESeq2 (alpha = 0.05). The RNA-Seq data have the accession no. E-MTAB-8166 and were uploaded on https://www.ebi.ac.uk/biostudies/arrayexpress/studies/E-MTAB-8166?key=64a67428-adb9-4681-99c9-98910b78ed4c. The genes differentially expressed between WT and Tau-KO cells (734) were used to interrogate a possible gene-set enrichment utilizing the transcription tool of the EnrichR portal (Chen et al., 2013, Kuleshov et al., 2016, Xie et al., 2021).

### 2.3 Pseudoviral particle production and transduction

Pseudolentiviral particles were produced by transient transfection of HEK293FT cells with 2 μg of the pSIF-H1-puro-IGFBP3 shRNA-2 or of the control plasmid pSIF-H1-puro-luciferase shRNA and 8 μg of the feline immunodeficiency virus (FIV) packaging plasmid mix (pFIV-34N & pVSV-G); all plasmids were kindly provided by Prof. Yuzuru Shiio (Greehey Children’s Cancer Research Institute, University of Texas). Cell conditioned medium was collected 2 days after transfection and cleared by centrifugation at 300 g for 5 min, 4°C. Pseudo-lentiviruses were 20-fold concentrated with centrifugal filters (MWCO 30 kDa, UFC903024, Amicon) at 3’000 g for 30-45 min, 4°C, aliquoted and stored at -80°C.

Human neuroblastoma SH-SY5Y cells (1 ×10^5^) were seeded into a 24-well plate coated with poly-D-lysine (p6407, Sigma-Aldrich) one day before pseudo-lentiviral particle transduction. One day after transduction, cells were supplemented with fresh complete medium and selected in the presence of 2.5 μg/mL puromycin (P8833, Sigma-Aldrich) for two weeks.

### 2.4 Drugs and cell treatments

Treatment of SH-SY5Y cells with Tazemetostat (CAS No. 1403254-99-8, S7128, Selleckchem) was performed at 10 µM for four days staring from a 10 mM stock solution in DMSO; vehicle 0.1% DMSO was added to the controls.

### 2.5 Western blot and immune precipitation

For direct analysis by western blot, total lysates from cells cultured in 6-well plates were prepared in 50 μL of SDS-PAGE sample buffer (1.5% SDS, 8.3% glycerol, 0.005% bromophenol blue, 1.6% β-mercaptoethanol and 62.5 mM Tris pH 6.8) and incubated for 10 min at 100°C. 15 μL per lane of the sample was loaded on SDS-polyacrylamide gels (SDS-PAGE).

For immune isolation, the cells were rinsed with PBS and lysed on ice in 100 μL of AlphaLisa Lysis Buffer (AL003, PerkinElmer) supplemented with protease and phosphatase inhibitor cocktails (S8820 & 04906845001, Sigma-Aldrich). Cell lysates were treated with benzonase (707463, Novagen) for 15 min at 37°C, centrifuged at 20’000 g for 10 min at 4°C and supernatants were collected as cell extracts. These latter were diluted in HiBlock buffer (10205589, PerkinElmer) and incubated overnight at 4°C with 0.5 μg of primary antibodies against SUZ12 (3737, Cell Signaling Technology), or EZH2 (5246, Cell Signaling Technology). Protein G-Sepharose beads (101241, Invitrogen) were added for 1 h at room temperature (RT) and the beads were washed three times in PBS with 0.1% Tween-20. Bead-bound proteins were eluted in SDS-PAGE sample buffer by boiling for 10 min at 100°C.

After SDS-PAGE, PVDF membranes with transferred proteins were incubated with primary antibodies: 0.084 μg/mL SUZ12, 0.421 μg/mL EZH2, 0.18 μg/mL GAPDH (ab181602, Abcam), 0.1 μg/mL H3K27me3 (C15410069, Diagenode), 0.02 μg/mL H3 (ab176842, Abcam), or 0.4 μg/mL IGFBP3 (sc365936, Santa Cruz Biotechnology). Primary antibodies were revealed with anti-mouse IgG coupled to IRDye RD 680 or anti-rabbit IgG coupled to IRDye 800CW (Licor Biosciences, 926– 68070 & 926–32211) on a dual infrared imaging scanner (Licor Biosciences, Odyssey CLx 9140) and quantified with the software provided (Licor Biosciences, Image Studio V5.0.21, 9140–500).

### 2.6 Immune staining

For immune staining, cells were grown on poly-D-lysine coated 8-well microscope slides (80826-IBI, Ibidi). Cells were fixed in 4% paraformaldehyde and stained (Ulrich et al., 2018) with primary antibodies: 0.168 μg/mL SUZ12, 0.842 μg/mL EZH2, 1.6 μg/mL H3K27me3 or 1.5 μg/mL p16 (ab108349, Abcam). Detection by fluorescent laser confocal microscopy (Nikon C2 microscope) was done with 2 μg/mL secondary antibodies anti-mouse IgG Alexa594, anti-rabbit IgG -Alexa 488 or anti-rabbit IgG-Alexa 647 (A-11032, A-11034, A21245, Thermo Fisher Scientific). Nuclei were counterstained with 0.5 μg/mL DAPI (D9542, Sigma-Aldrich). Images were acquired by sequential excitations (line-by-line scan) with the 405 nm laser (464/40 emission filter), the 488 nm laser (525/50 nm filter), the 561 nm laser (561/LP nm filter) and the 650 nm laser (594/633 emission filter). ImageJ was used for all image quantifications.

### 2.7 RNA extraction and RT-qPCR

Total RNA extraction using the TRIzol™ Reagent (15596026, Invitrogen) and cDNA synthesis using the GoScript Reverse Transcription Mix Random Primers (A2800, Promega) were done according to the instructions of the manufacturer. Amplification was performed with SsoAdvanced Universal SYBR Green Supermix (1725271, BioRad) with 43 cycles at 95°C for 5 sec, 60°C for 30 sec and 60°C for 1 min using specific primers for *EZH2, SUZ12, IGFBP3, GPR37, ITGA3, MRC2* and *IRF6* gene transcripts **(Supplementary Table IV)**. Relative RNA expression was calculated using the comparative Ct method and normalized to the geometric mean of the GAPDH and HPRT1 mRNAs.

### 2.8 SA-βGal assay

Senescence-associated β-galactosidase (SA-βGal) staining was determined on cells grown in 6-well plates, fixed with 2% paraformaldehyde for 10 min at RT and washed twice with gentle shaking for 5 min at RT. Cells were then incubated with 1 mg/mL X-gal (20 mg/mL stock in DMF; B4252, Sigma-Aldrich,) diluted in pre-warmed 5 mM K_3_[Fe(CN)_6_] (P-8131, Sigma-Aldrich), 5 mM K_4_[Fe(CN)_6_]·3H_2_O (P-3289, Sigma-Aldrich), and 2 mM MgCl_2_ (M-8266, Sigma-Aldrich) in PBS at pH 6.0. Acquisition and quantification of the images for SA-βGal activity and cell area were done with an automated live cell imager (Lionheart FX, BioTek).

### 2.9 LysoTracker

For LysoTracker staining, cells were seeded in poly-D-lysine coated 8-well microscope slides, incubated for 10 min at 37°C with 0.25 μM Lysotracker Red (L7528, Thermo Fischer Scientific). Nuclei were counterstained with 2.5 μg/mL Hoechst (H3570, Invitrogen) for 10 min at 37°C, afterwards cells were washed with complete medium. Images of living cells were acquired on a fluorescent laser confocal microscope (C2, Nikon) by sequential excitations (line-by-line scan) with the 405 nm laser (464/40 nm emission filter), and the 561 nm laser (561/LP nm filter). ImageJ was used for all image quantifications. Both the lysosomes area and the mean number of lysosomes per cells and per images were determined.

## 3 Results

### 3.1 Transcriptomics analysis of Tau-KO cells

We performed a next-generation transcription (RNAseq) analysis of human SH-SY5Y neuroblastoma cells knocked-out for Tau when compared to normal Tau-expressing cells (**Fig 1A**). The sequences obtained from the six samples analyzed (three Tau expressing cell lines, three Tau-KO cell lines) mapped reliably to ∼16’000 genes. Additional 14’000 transcripts were expressed at low levels and were not included in the differential expression analysis. The primary data were stored at https://www.ebi.ac.uk/biostudies/ with open access (E-MTAB-8166). When filtering for differentially expressed transcripts in Tau-KO cells, 1388 RNAs displayed a significant change (Adj P <.05), of which 723 RNAs were up-regulated in the log2(FC) range between 0.31 and 11.05 (between 1.24 and ∼2000 fold higher than in control Tau expressing cells) (**Supplementary Table I**).

**Figure 1.**
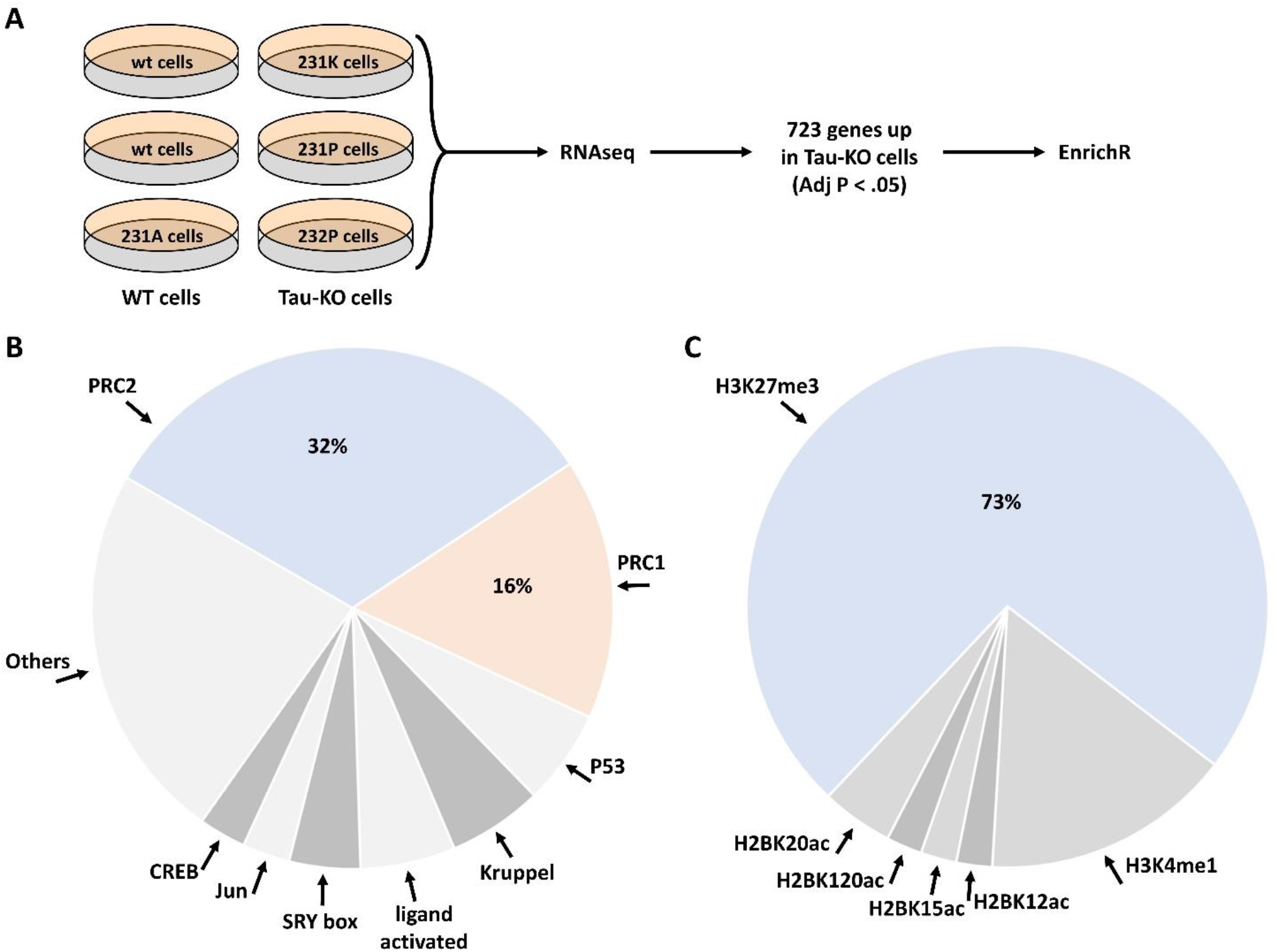
Deregulation of the PRC2 pathway in human Tau-KO SH-SY5Y neuroblastoma cells. (**A**). Scheme of the procedure for the RNAseq and EnrichR analyses in Tau-expressing (WT) and Tau-knock-out (Tau-KO) cells. (**B-C**). The EnrichR analysis based on 723 upregulated genes in Tau-KO cells resulted in the enrichment of the PRC2 pathway with the ChIP datasets (**B**) and of H3K27me3 with the epigenomics datasets (**C**).

We selected these 723 differentially expressed genes to interrogate a possible gene-set enrichment utilizing the transcription tool of the EnrichR portal (Chen et al., 2013, Kuleshov et al., 2016, Xie et al., 2021). The CHIP-sequencing datasets (ChEA 2016) identified an overrepresentation of Polycomb Repressive Complex-associated proteins among the 68 datasets showing a significant (Adj P <.01) difference. Indeed, almost half of the enriched CHIP-sequencing datasets were obtained from core components or known regulators of PRC2 (32%) or of PRC1 (16%) (**Fig 1B**; **Supplementary Table II**). PRC2 actively catalyzes the trimethylation of histone 3 (H3) at lysin 27 (H3K27me3) (Laugesen et al., 2016, Moritz and Trievel, 2018). In agreement with the identification of PRC2 in the CHIP-sequencing datasets, mining of the epigenomics roadmap (HM ChIP-seq) resulted in a 73% enrichment of the H3K27me3 signature (Adj P <.01) in 45 datasets (**Fig 1C**; **Supplementary Table III**). Altogether, analysis of the RNAseq data suggested that up-regulation of transcription of a specific set of genes in Tau-depleted neuroblastoma SH-SY5Y cells might ensue from a relief of gene activity decline controlled by PRC2.

### 3.2 Reduced expression and activity of PRC2 in Tau-depleted cell

We subsequently analyzed the amount of two PRC2 core components in Tau-KO cells by western blot. In agreement with what suggested by the transcriptomics data, we observed reduced amounts of the catalytic subunit EZH2 and the scaffold subunit SUZ12 of PRC2 in Tau-KO cells when compared to Tau-expressing cells (**Fig 2A**). Reduced proteins were found also by quantitative immune staining of the cells utilizing specific antibodies (**Fig 2B**). RT-qPCR analysis excluded that the effect on PRC2 protein resulted from reduced transcription since no difference was found for the EZH2 and SUZ12 mRNAs (**Fig 2C**), data that suggested a Tau-dependent effect on PRC2 protein stability. Nonetheless, co-immune isolation revealed the presence of the EZH2-SUZ2 core complex of PRC2 in Tau-KO cells, albeit at reduced levels when compared to controls (**Fig 2D**).

**Figure 2.**
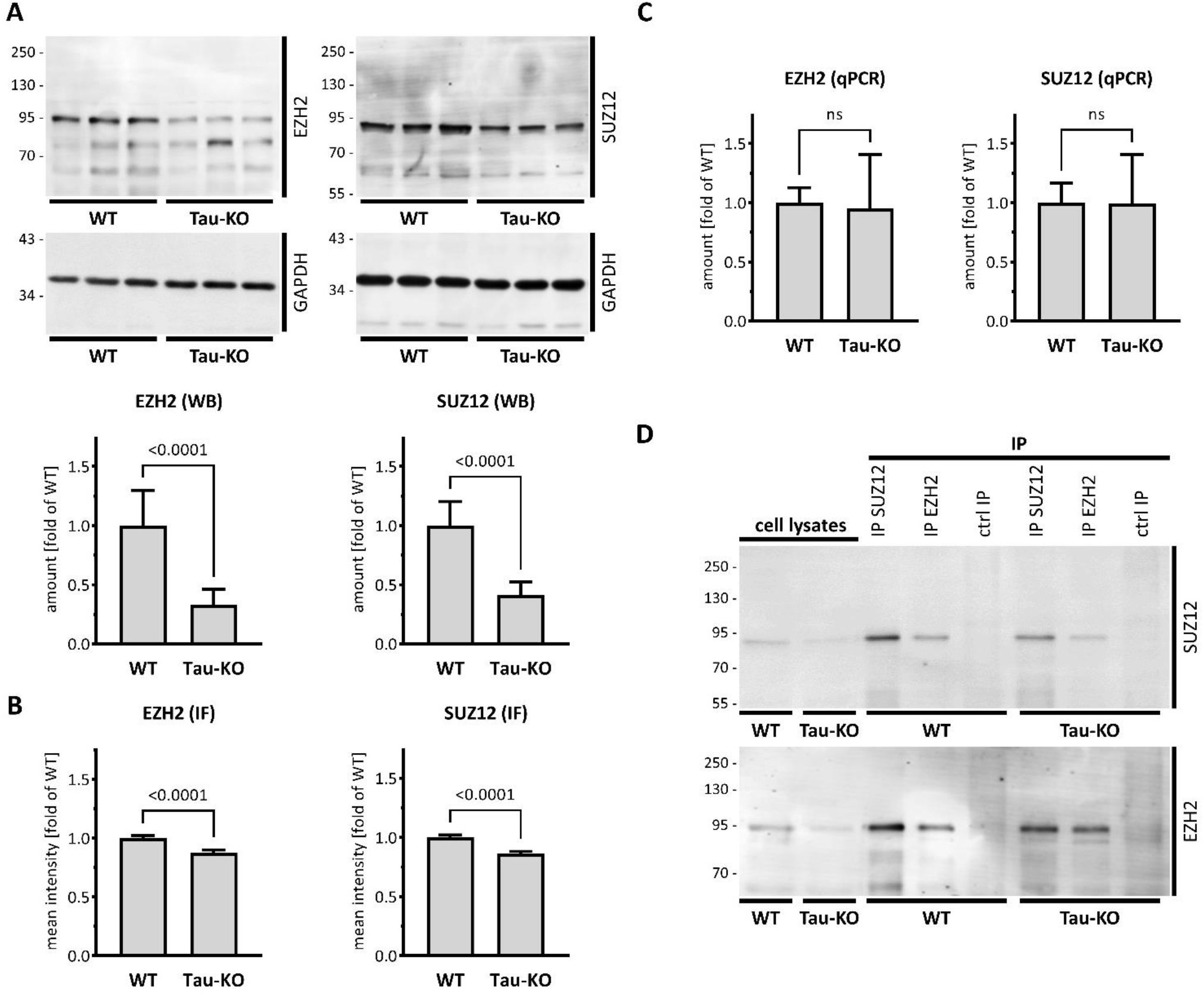
Reduced PRC2 complex in Tau-KO SH-SY5Y cells. (**A**). Shown are matched protein amounts of parental (WT) or Tau-KO cell lysates analyzed by western blot (biological triplicates on a single gel) with EZH2, SUZ12 or GAPDH primary antibodies and anti-rabbit IgG IRDye 800CW secondary antibody. The EZH2 and SUZ12 signals were normalized on the respective GAPDH signals and reported as fold of WT; mean ± SD of 9 biological replicates, unpaired Mann-Whitney test. (**B**). Determination of nuclear EZH2 or SUZ12 mean fluorescent intensity analyzed by immune fluorescence staining and laser confocal microscopy with EZH2 or SUZ12 antibodies revealed with an anti-rabbit AlexaFluor 488 antibody. Data obtained with a DAPI nuclear mask (ImageJ) are reported as fold of WT; mean ± SEM of 292-475 nuclei from two (EZH2) or three (SUZ12) independent experiments, unpaired Mann-Whitney test. (**C**). Shown are RT-PCR determination of mRNA with specific primers for EZH2 or SUZ12. Normalization was performed on the geometric mean of the GAPDH and HPRT1 mRNA values and reported as fold of WT; mean ± SD of 12 biological replicates, unpaired Mann-Whitney test. (**D**). Shown are matched protein amounts of cell lysates subjected to immune isolation (IP) with EZH2 or SUZ12 antibodies or matched amounts of control antibodies (ctrl IP). Samples were resolved on a single same gel and analyzed by western blot with EZH2 or SUZ12 antibodies and secondary anti-rabbit IgG IRDye 800CW antibody.

PRC2 enzymatic activity was determined by western blot and quantitative immune staining of H3K27me3, an epigenetic mark produced by the histone methyl transferase activity of PRC2 (Guo et al., 2021). Confirming the lower amounts of the PRC2 complex, we found that Tau-KO cells display reduced H3K27me3 (**Fig 3A**-**B**). Among the upregulated transcripts found by RNA-seq in Tau-KO cells, we selected five known PRC2 targets displaying close to average signals (**Supplementary Table I**): *IGFBP3* (19.0x of WT, adjP .004), *GPR37* (9.3x, .015), *ITGA3* (6.3x, .017.3x), *MRC2* (5.1x, .016), and *IRF6* (3.2x, .0498). Determination of mRNA expression by RT-qPCR validated their up-regulation (**Fig 3C**); indicating again that Tau-depletion relieved repression of transcription caused by reduced PRC2 activity in Tau-depleted cells.

**Figure 3.**
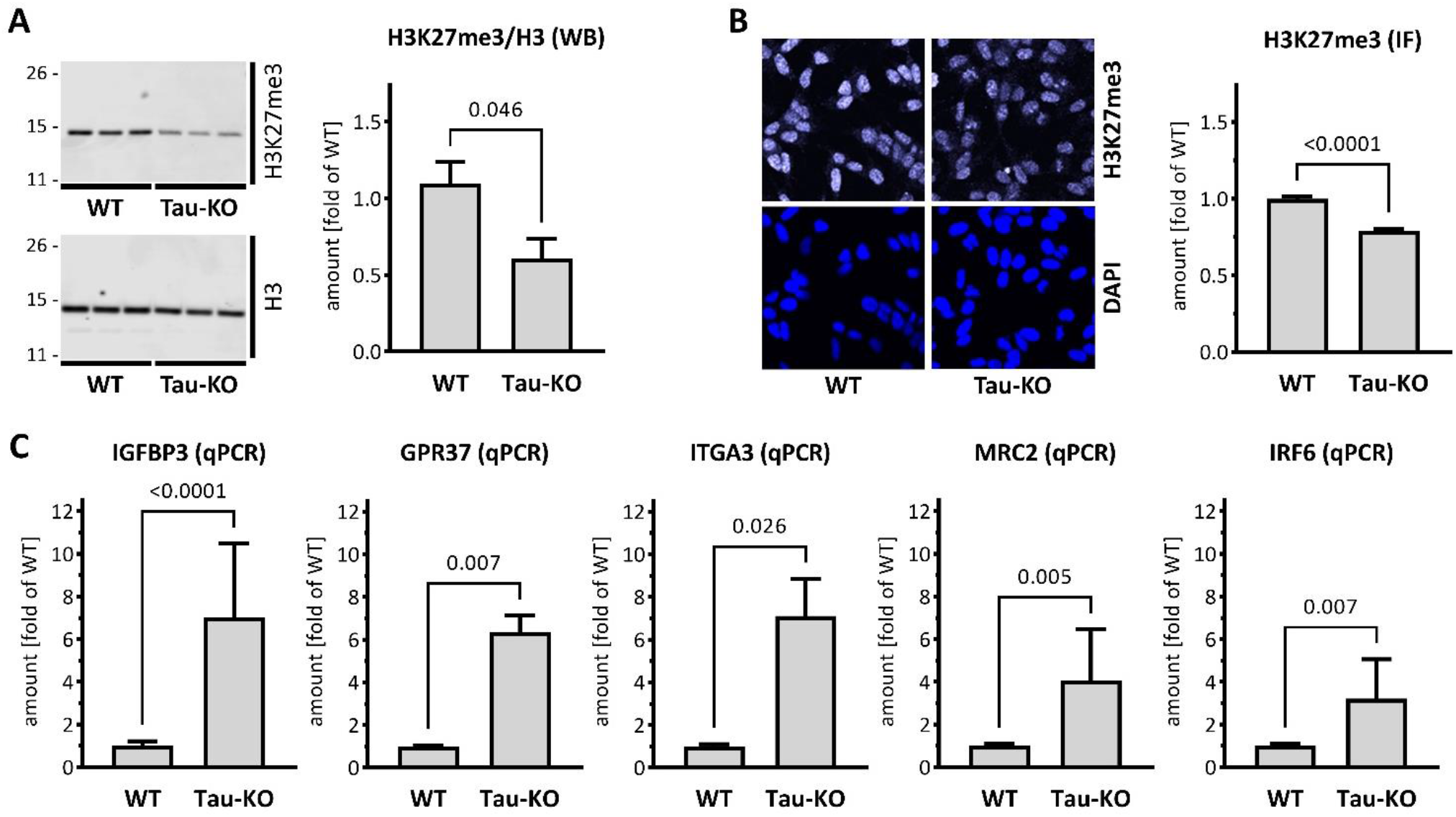
Reduced PRC2 activity in Tau-KO SH-SY5Y cells. (**A**). Shown are matched protein amounts of parental (WT) or Tau-KO cell lysates analyzed by western blot (biological triplicates on a single gel) with H3K27me3, H3 or GAPDH primary antibodies and anti-rabbit IgG IRDye 800CW secondary antibody. The H3K27me3 and H3 signals were normalized for GAPDH and reported as fold of WT for the H3K27me3/H3 ratios; mean ± SD of 8-9 biological replicates, unpaired Mann-Whitney test. (**B**). Nuclear H3K27me3 mean fluorescent intensity was determined by immune fluorescence staining and laser confocal microscopy. Data obtained with a DAPI nuclear mask (ImageJ) are reported as fold of WT; mean ± SEM of 695-716 nuclei from five independent experiments, unpaired Mann-Whitney test. (**C**). Shown are RT-PCR determination of mRNA with specific primers as indicated. Normalization was performed on the geometric mean of the GAPDH and HPRT1 mRNA values and reported as fold of WT; mean ± SD of 3-12 biological replicates, unpaired Mann-Whitney test.

### 3.3 PRC2-dependent overproduction of IGFBP3 in Tau-KO cells

IGFBP3 protein is a component of the senescence-associated secreted phenotype (SASP) (Basisty et al., 2020). Having previously reported that Tau-depletion favored cellular senescence (Sola et al., 2020), we interrogated the role of IGFBP3 in this process. As anticipated from the mRNA data, a strong overproduction of endogenous IGFBP3 was present in Tau-KO cells (**Fig 4A**). Reinforcing the link between PRC2 and IGFBP3, treatment of SH-SY5Y cells with Tazemetostat, a specific blocker of the histone methyl transferase activity of EZH2 (Straining and Eighmy, 2022), reduced H3K27me3 and increased IGFBP3 (**Fig 4B**). Lower EZH2 and SUZ12 and higher IGFBP3 was confirmed in an independent Tau-KO cell line (**Supplementary Fig 1**).

**Figure 4.**
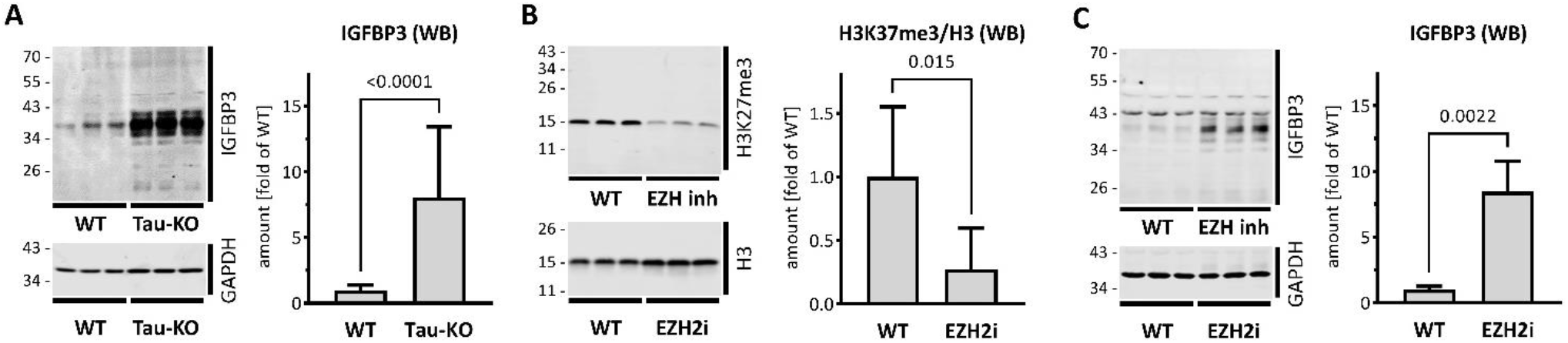
Increased IGFBP3 in SH-SY5Y cells after Tau-KO or PRC2 inhibition. (**A**). Shown are matched protein amounts of parental (WT) or Tau-KO cell lysates analyzed by western blot (biological triplicates on a single gel) with IGFBP3 or GAPDH primary antibodies and anti-mouse IgG IRDye 680RD or anti-rabbit IgG IRDye 800CW secondary antibodies. The IGFBP3 signals were normalized for GAPDH and reported as fold of WT; mean ± SD of 8-9 biological replicates, unpaired Mann-Whitney test. (**B-C**). Shown is a western blot of matched protein amounts of SH-SY5Y cells treated for 4 days in the absence (WT) or presence of 10 μM Tazemetostat (EZH2i). Biological triplicates on a single gel were probed (**B**) with H3K27me3, H3 or GAPDH primary antibodies and anti-rabbit IgG IRDye 800CW secondary antibody or (**C**) with IGFBP3 or GAPDH antibodies. Protein signals were normalized for GAPDH and reported as fold of WT; mean ± SD of 6 biological replicates, unpaired Mann-Whitney test.

### 3.4 Tau/PRC2/IGFBP3 triad in senescence

Increased cellular expression of IGFBP3 is associated with autocrine and paracrine senescence induction (Elzi et al., 2012), and reduced PRC2 is also linked to increased cellular senescence (Ito et al., 2018). Furthermore, increased senescence is observed in Tau-KO cells (Sola et al., 2020). Thus, we postulated that PRC2-dependent de-repression of IGFBP3 in Tau-depleted cells may explain the induction of cellular senescence. We observed first that Tau depletion as well as PRC2 inhibition both increased the percentage of SH-SY5Y cells entering in senescence, as assessed by three independent markers: P16, senescence-associated β-galactosidase (SA-βgal), and the number and size of lysosomes labelled with the acidotrophic LysoTracker dye (**Fig 5A**). Next, we reduced IGFBP3 expression in Tau-KO cells by a shRNA-based approach **(Fig 5B**) and found that this reduced senescence induction in Tau-KO cells **(Fig 5C)**. Thus, we validated our hypothesis that increased senescence in Tau-KO SH-SY5Y cells was likely the consequence of decreased PRC2-dependent repression of IGFBP3 expression.

**Figure 5.**
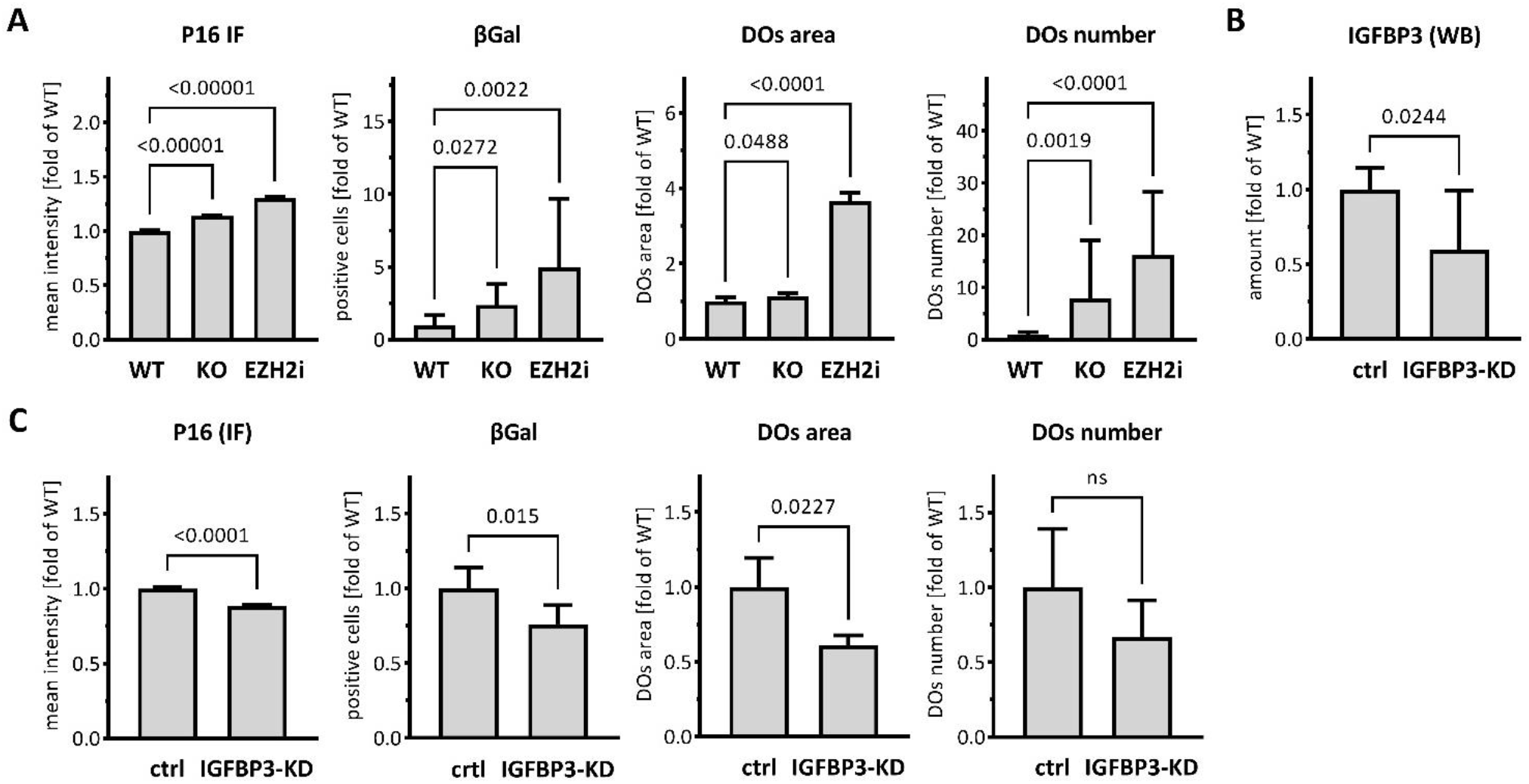
Increased IGFBP3-induced senescence in cells. (**A**). Senescence markers were analyzed in parental (WT), Tau-KO or 10 μM Tazemetostat-treated parental (EZH2i) cells. The nuclear P16 fluorescent intensity (P16 IF) was determined with a DAPI nuclear mask (ImageJ) by immune fluorescence staining and laser confocal microscopy and reported as fold of WT; mean ± sem of 1991-3153 nuclei. The percent of senescence-associated βGal positive cells (βGal) was determined by automated cell imaging; mean ± SD of 7-14 fields. Living cells labeled with the acidotrophic dye LysoTracker (DOs) were imaged on a laser confocal microscope and analyzed for the size (area of 643-3429 DOs) and mean number per cell (13-18 fields) of LysoTracker-positive organelles (ImageJ). Values are reported as fold of WT; mean ± sem (area) or ± SD (number), non-parametric Krustal-Wallis and Dunn’s multiple comparison test. (**B**). Matched protein amounts of SH-SY5Y cells transduced with mock (ctrl) or IGFBP3 shRNA (IGFBP3-KD) pseudo lentiviral particles were analyzed by western blot (biological triplicates on a single gel) with IGFBP3 or GAPDH primary antibodies and anti-mouse IgG IRDye 680RD or anti-rabbit IgG IRDye 800CW secondary antibodies. The IGFBP3 signals were normalized for GAPDH and reported as fold of WT; mean ± SD of 9 biological replicates, unpaired Mann-Whitney test. (**C**). As in (**A**) for SH-SY5Y cells transduced with mock (ctrl) or IGFBP3 shRNA (IGFBP3-KD) pseudo lentiviral particles. P16 IF: mean ± sem of 669-881 nuclei; βGal: mean ± SD of 6 fields. DOs; area of 456-551 DOs, mean number per cell (5 fields) of LysoTracker-positive organelles (ImageJ). Values are reported as fold of WT; mean ± sem (area) or ± SD (number); unpaired Mann-Whitney test.

## 4 Discussion

We report data demonstrating a non-canonical role of Tau as a modulator of the epigenetic activity of PRC2 inducing cellular senescence in neuroblastoma SH-SY5Y cells. In our study we identified a prevalent PRC2 signature for shared modulation of upregulated transcripts in Tau-depleted cells. PRC1 and PRC2 are multi-subunit transcriptional repressors that crucially modulate chromatin structure and gene expression by distinct enzymatic activities (Vijayanathan et al., 2022). PRC1 is an E3 ubiquitin ligase that catalyzes H2A ubiquitination at lysine119, whereas PRC2 acts as a methyltransferase that generates H3K27me3 with some cross-talk between the two complexes (Guerard-Millet et al., 2021). Confirming the bioinformatics results, Tau depletion caused reduced cellular amounts of PRC2 and its product H3K27me3. Increased senescence status upon Tau-depletion was reproduced through pharmacological inhibition of PRC2 in Tau-expressing cells. Finally, we report that reversing the up-regulation of the PRC2-target IGFBP3, impaired senescence induction in Tau-depleted cells.

Evidence exists for the implication of Tau in chromatin remodeling. A pioneering study investigated chromatin conformation in mouse and drosophila models of AD as well as in human diseased brain, whereby a general loss of heterochromatin was associated with aberrant gene expression in all three paradigms (Frost et al., 2014). More recently, binding of Tau to histones was linked to the maintenance of condensed chromatin (Rico et al., 2021). Thus, Tau may favor chromatin compaction for preventing aberrant gene transcription. Misfolding, hyperphosphorylation or sequestration of Tau in oligomers and fibrils, typical hallmarks of tauopathies, could all result in a negative regulation of this non-canonical function of Tau. In our study, we show that in addition to the direct interaction between Tau and histones (Rico et al., 2021), an indirect mechanism involving PRC2 is an additional instrument for modulating chromatin compaction.

The role of PRC2 in senescence was shown by findings indicating that impairment of its catalytic activity induces a delayed decrease in H3K27me3 at the CDKN2A locus, which upregulates p16, the SASP phenotype, and senescence (Ito et al., 2018). This function of PRC2 represents a target for anticancer therapies e.g., through EED inhibition associated to de-repression of SASP-encoding genes and entry of proliferative cancer cells in a senescent state (Chu et al., 2022). Beside the paradoxical implication in cancer (Yang et al., 2021), senescence contributes to neurodegenerative diseases. Senescent neurons, microglia, astrocytes and neuronal stem cells were found during the pathogenic process (Si et al., 2021). A recent study in a tauopathy mouse model supported a causal link between cell senescence and cognitive decline linked to neuronal loss. Indeed, p16INK4A-positive senescent glial cells were found associated to Tau lesions, and, strikingly, the clearance of these cells prevented Tau hyperphosphorylation, Tau fibril deposition, whilst preserving neuronal survival and cognitive functions (Bussian et al., 2018). We describe now a conceivable mechanism linking depletion of functional Tau in tauopathies and senescence induction.

Among the PRC2 targets, we identified IGFBP3 as a main driver of senescence resulting from Tau-depletion. Ectopic expression of the SASP component IGFBP3 or its administration to MCF7 or IMR-90 cells is sufficient to induce senescence, whereas IGFBP3 knock-down impairs doxorubicin-induced senescence (Elzi et al., 2012). Although the role of PRC2 and IGFBP3 were established independently, to our knowledge our study is the first one showing IGFBP3 as a main executor of PRC2-dependent senescence induction and its modulation by Tau protein levels.

PRC2 has numerous functions in the developing central nervous system, with many neurogenesis-linked genes regulated by the PRC2/H3K27 axis (Liu et al., 2017). PRC2 is essential in preserving neural progenitor cell identity and neuroepithelial integrity (Akizu et al., 2016). PRC2 deficiency in mice leads to aberrant gastrulation and lack of neural tissue (Schumacher et al., 1996). Later in development, a transcription pattern with a PRC2 signature drives neuronal migration and is essential for the organization of neural circuits (Zhao et al., 2015). The rare Weaver syndrome linked to developmental cognitive deficits is caused by autosomal dominant mutations in any one of the three PRC2 core components EZH2, EED and SUZ12 (Deevy and Bracken, 2019). However, PRC2 is also involved in neurodegeneration. PRC2 deficiency in striatal neurons of mice reactivates the deleterious expression of transcripts that are normally suppressed in these cells, ultimately causing premature lethality (Von Schimmelmann et al., 2016). Additional studies implicated PRC2 in ataxia-telangiectasia (Li et al., 2013), Parkinson’s disease, Huntington’s disease and AD (Kuehner and Yao, 2019). A meta-analysis of differentially methylated regions in prefrontal neocortex at different disease stages has identified in AD several hypermethylated regions, which were significantly enriched in polycomb repressed regions (Zhang et al., 2020). These data also link PCR2-dependent methylation of H3 with that of CpG islands of the genome, another epigenetic mechanism of gene repression (Phillips, 2008).

PRC2 lacks sequence-specific DNA-binding ability and therefore relies on accessory proteins for targeting specific loci. Factors contributing to selective PRC2 recruitment to chromatin are the interaction with sequence-specific transcription factors or RNAs, and or discerning chromatin features (Blackledge and Klose, 2021). In the fly, PRC2 activity is regulated through the interaction with transcription factors binding to polycomb response elements often located in proximal promoter regions of developmental genes (Kassis and Brown, 2013). However, the orthologue system was not found in mammals (Bauer et al., 2016). Rather, it is maybe replaced by the evolution of a mechanism based on non-methylated CpG islands (Ku et al., 2008) and the action of DNA-binding proteins binding to them. Proteins with such features are the PRC1.1 complex member KDM2B, or the PRC2-members PHF1, MTF2 and PHF19 (Owen and Davidovich, 2022).

PRC histone modifications are heritable over mitotic cell division providing an epigenetic memory for stable cell identity and adequate response to stress (Reinig et al., 2020). Thus, PRC2 dysfunction is frequently associated with neoplastic progression and is a target for anticancer therapy (Comet et al., 2016). Expression of its catalytic subunit EZH2 correlates with cell proliferation, and its aberrant overexpression is frequent in many types of cancer cells (Liu and Liu, 2022). However, in line with a role in tumor suppression, loss-of-function of PRC2 is also involved in cancer (Liu et al., 2017). PRC2 modulation by Tau implicates this latter in the pathogenesis of cancer, supporting the observation that the Tau mRNA correlates with survival in several tumors (Gargini et al., 2019, Papin and Paganetti, 2020). The mechanism explaining this correlation is unknown but may involve the non-canonical role of Tau in modulating chromatin compaction and senescence induction. This may open new therapeutic opportunities for neurodegenerative diseases and cancer.

## 6 Author Contributions

Conceptualization: SP, PP

Methods and investigations: CM, MS, EP. MB, AR

Supervision: SP, PP

Writing (original draft): SP

Writing (review & editing): all co-authors

## 8 Acknowledgments

We thank the whole laboratory for support and advice during this study.

## 9 Data Availability Statement

The RNA-Seq data have the accession no. E-MTAB-8166 and were uploaded on https://www.ebi.ac.uk/biostudies/arrayexpress/studies/E-MTAB-8166?key=64a67428-adb9-4681-99c9-98910b78ed4c.

---

**Supplementary Figure 1.**
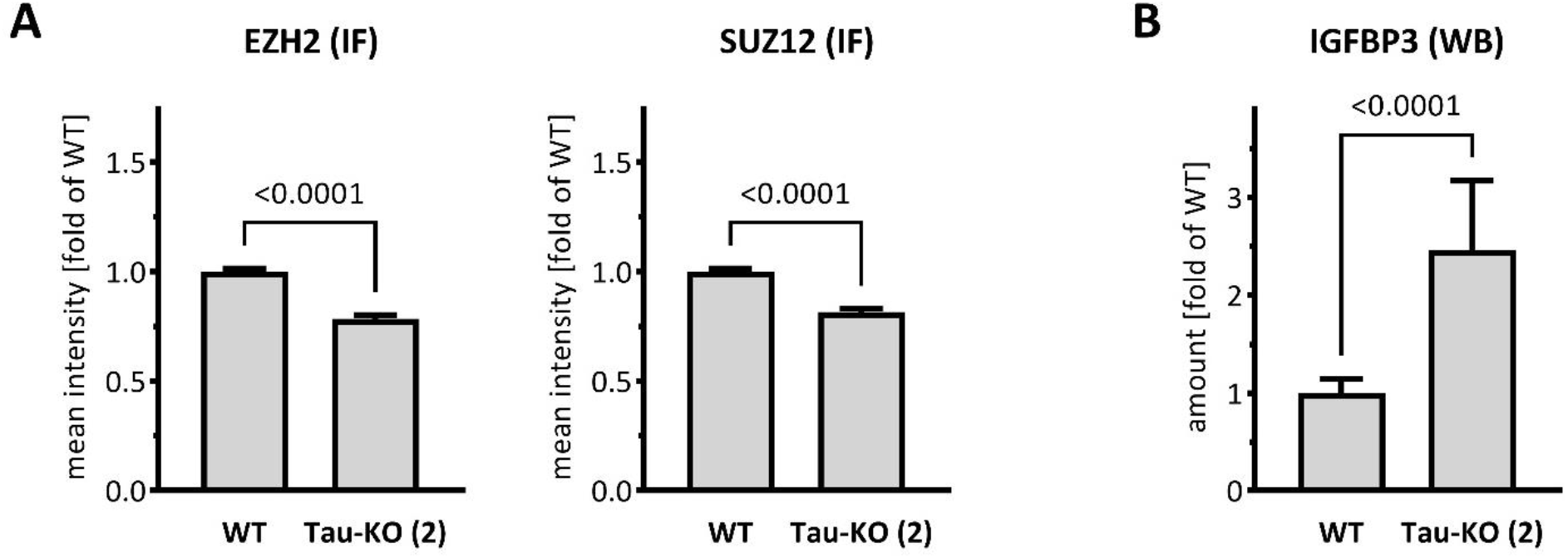
Reduced EZH2/SUZ12 and increased IGFBP3 in an independent Tau-KO SH-SY5Y cell line. (**A**). Nuclear EZH2 or SUZ12 mean fluorescent intensity was analyzed by immune fluorescence staining and laser confocal microscopy with EZH2 or SUZ12 antibodies revealed with an anti-rabbit AlexaFluor 488 antibody. Data obtained with a DAPI nuclear mask (ImageJ) are reported as fold of WT; mean ± sem of 202-208 nuclei (EZH2) or 182-184 nuclei (SUZ12), unpaired Mann-Whitney test. (**B**). IGFBP3 was determined by western blot for matched protein amounts of parental (WT) or Tau-KO (2) cell lysates (biological triplicates on a single gel). Data are reported as fold of WT, mean ± SD of 9 biological replicates, unpaired Mann-Whitney test.

**Supplementary Table I.**
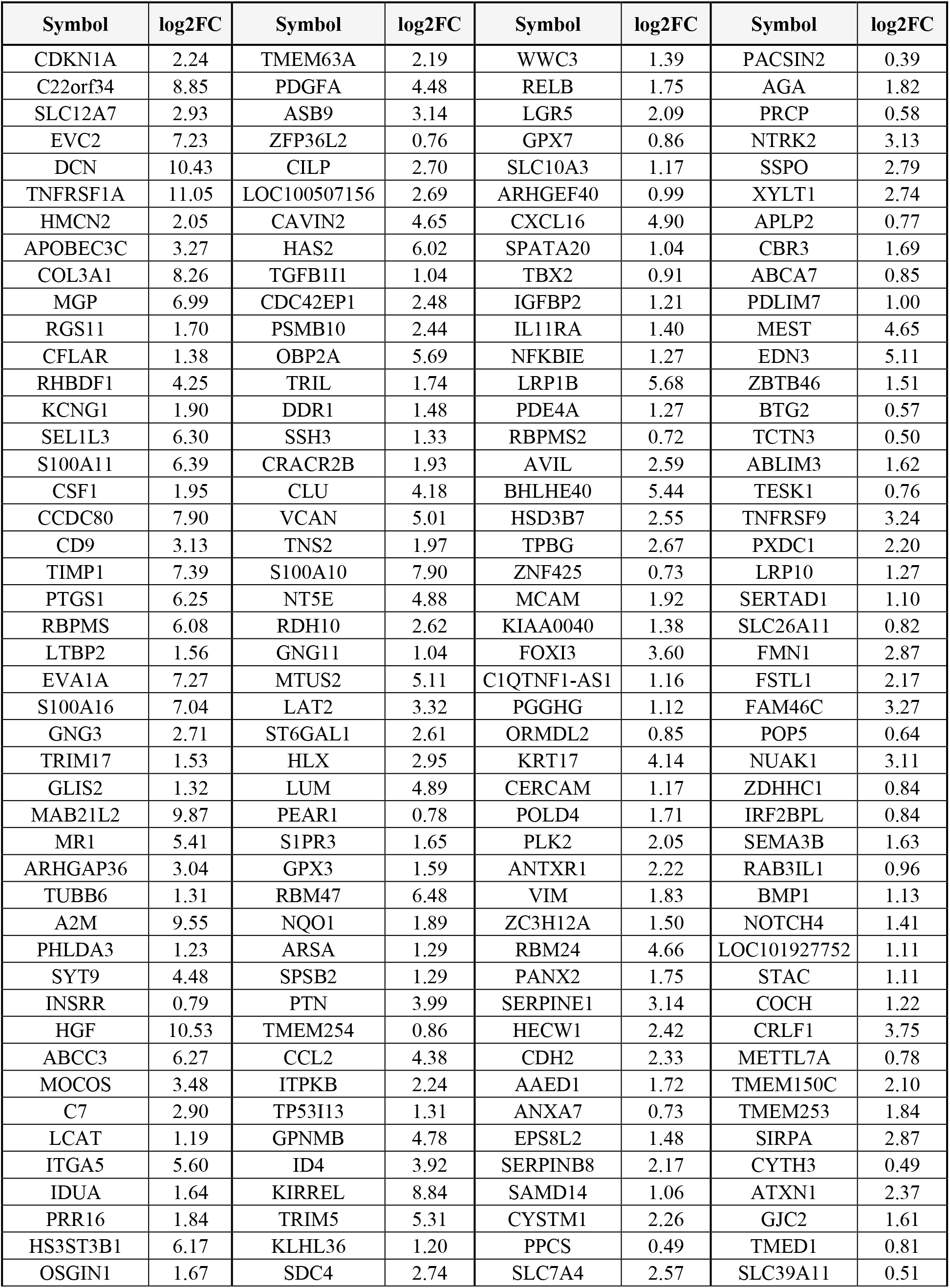
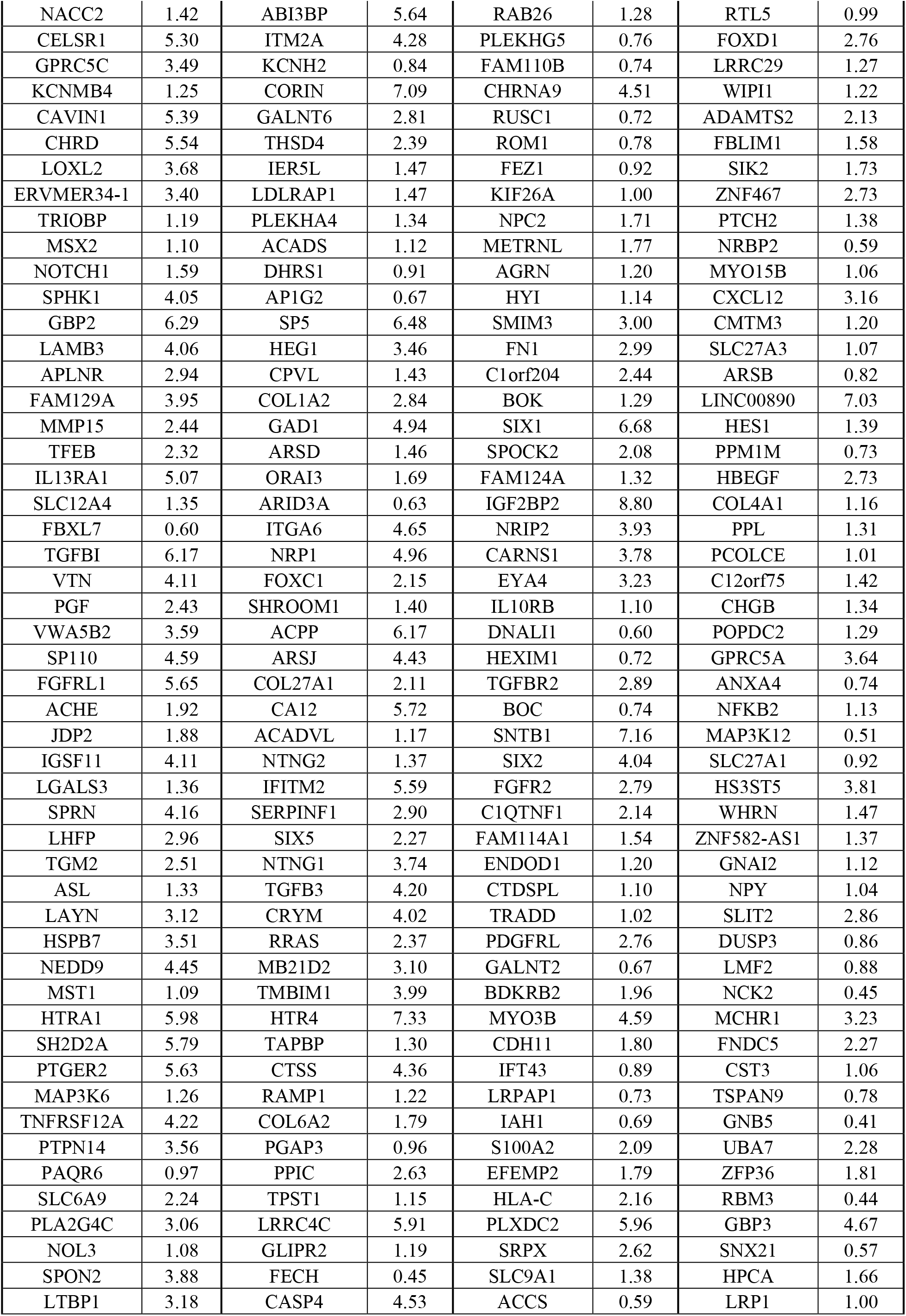
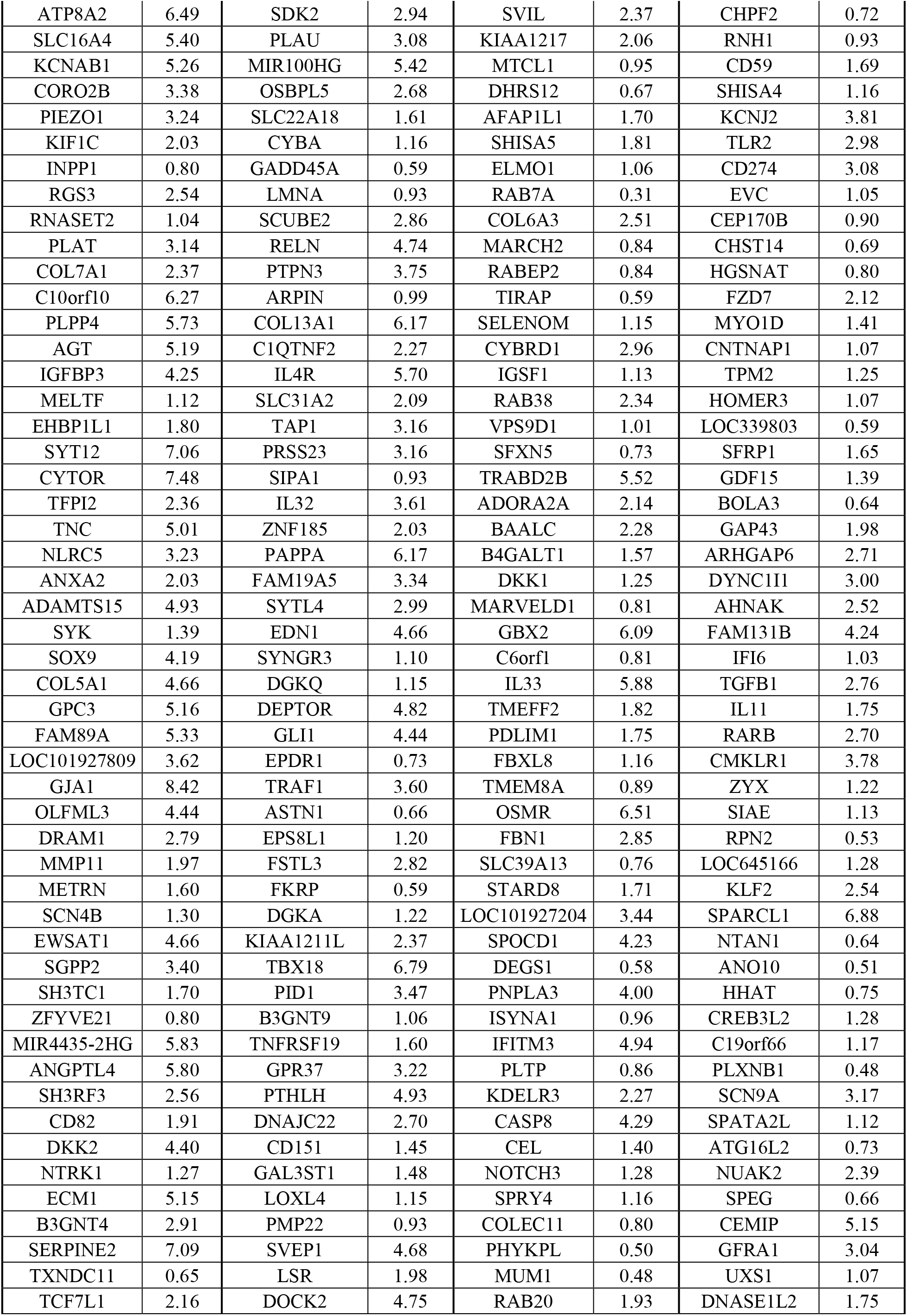
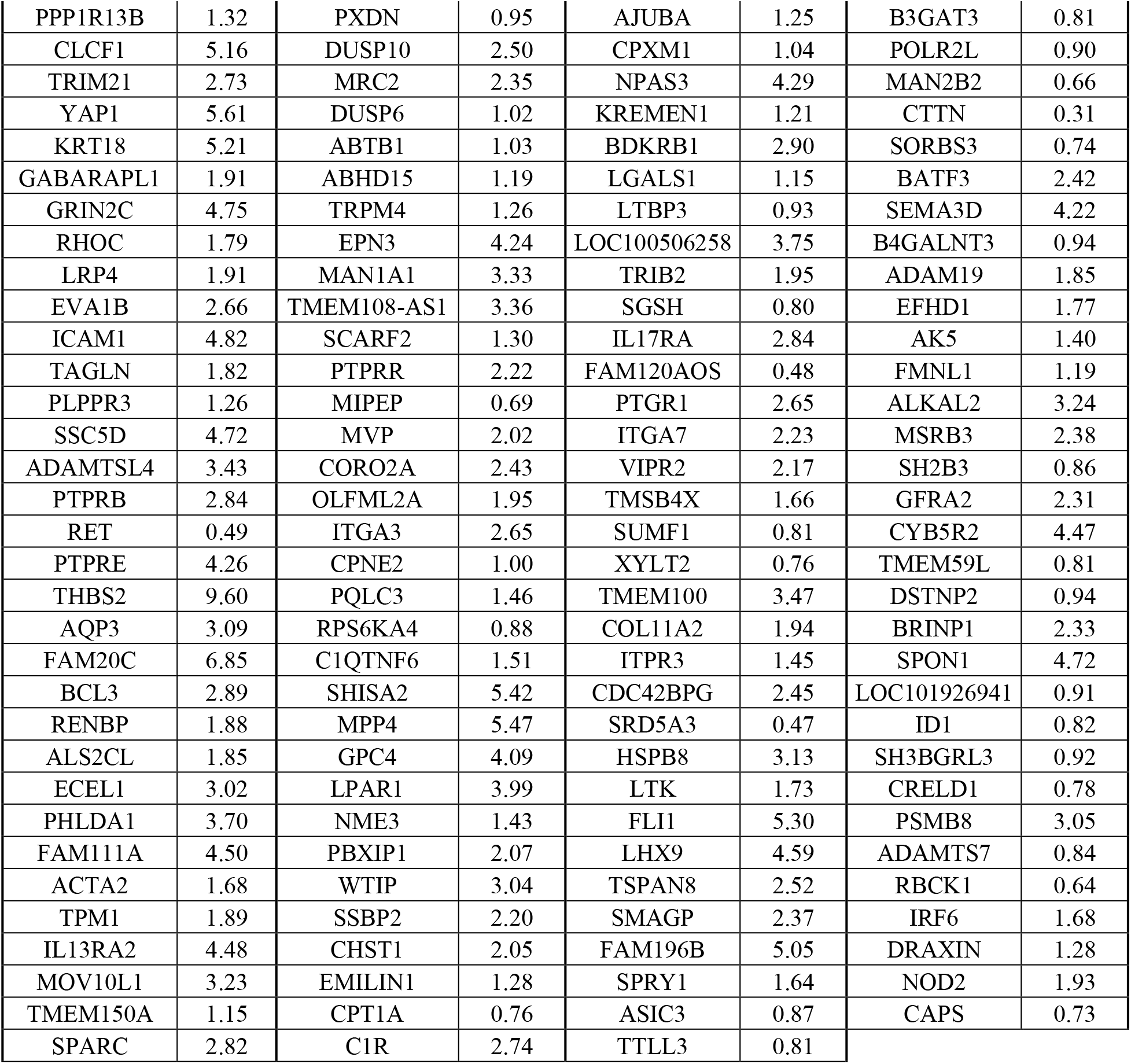
Transcripts upregulated in human Tau-KO SH-SY5Y cells (Adj P <.05) listed in order of decreasing significance.

**Supplementary Table II.**
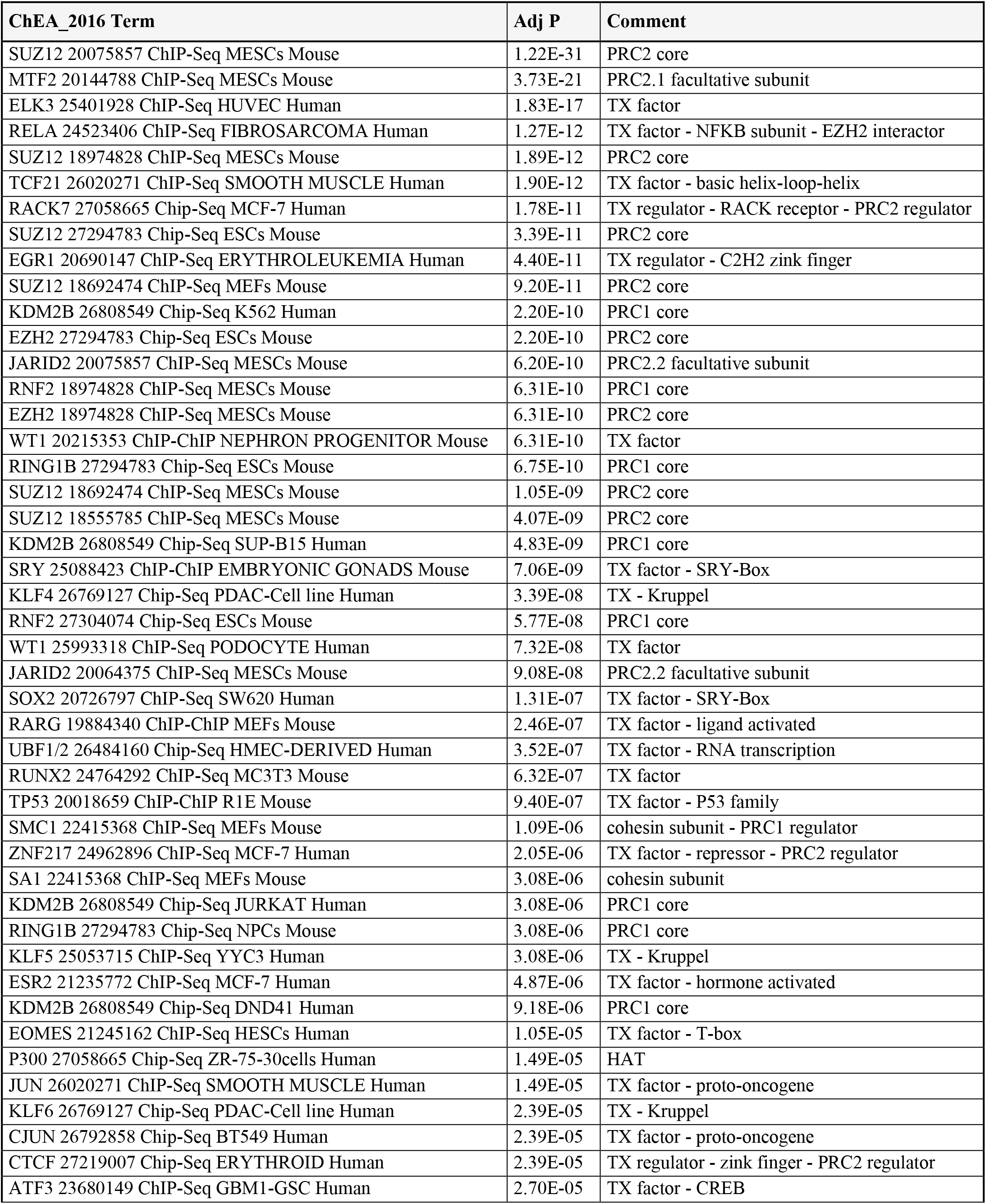
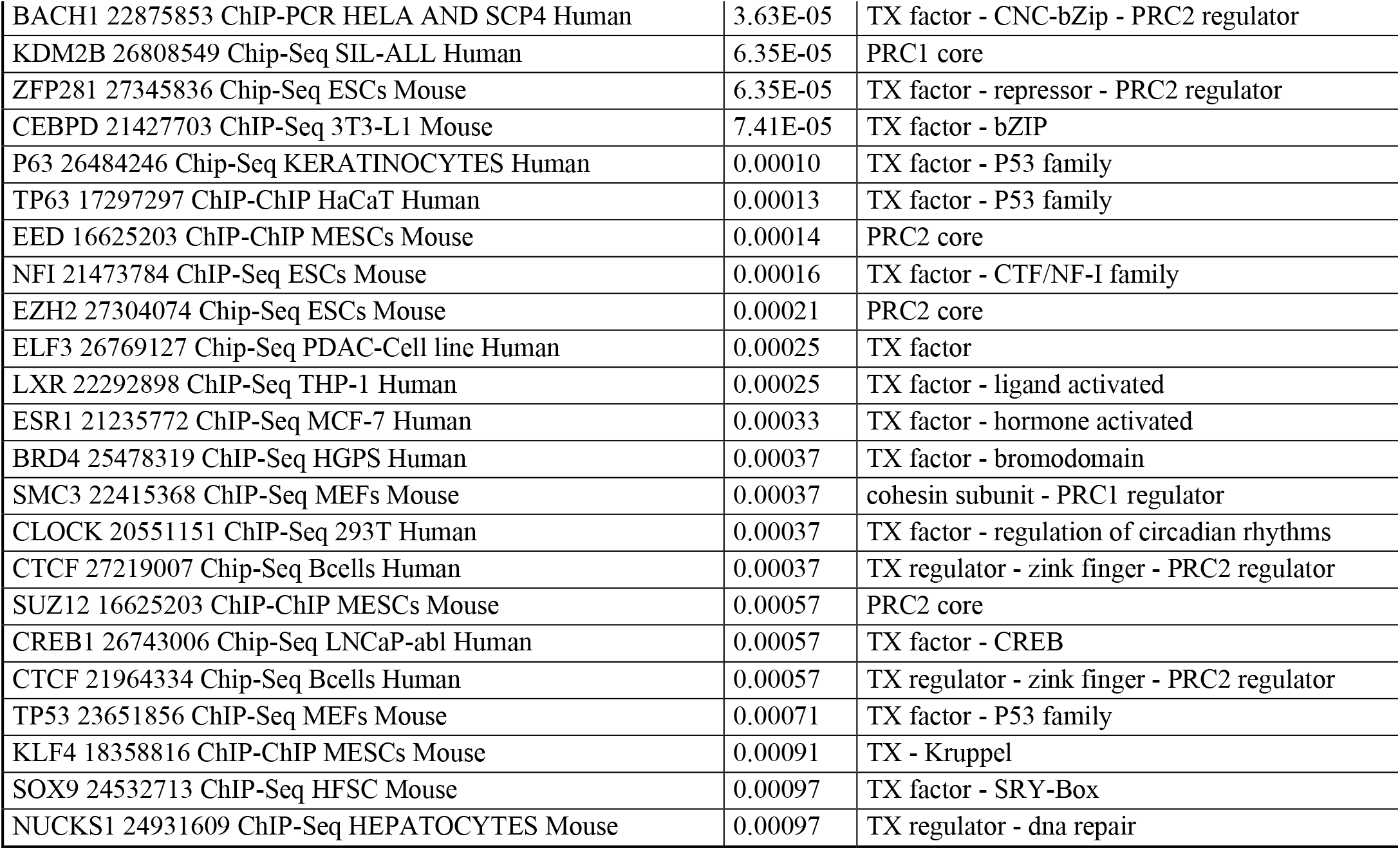
CHiP datasets identified with 723 upregulated transcripts in human Tau-KO cells (Adj P<.01). Analysis performed August 4^th^ 2022.

| ChEA_2016 Term | Adj P | Comment |
| --- | --- | --- |
| SUZ12 20075857 ChIP-Seq MESC's Mouse | 1.22E-31 | PRC2 core |
| MTF2 20144788 ChIP-Seq MESC's Mouse | 3.73E-21 | PRC2.1 facultative subunit |
| ELK3 25401928 ChIP-Seq HUVEC Human | 1.83E-17 | TX factor |
| RELA 24523406 ChIP-Seq FIBROSARCOMA Human | 1.27E-12 | TX factor - NFkB subunit - EZH2 interactor |
| SUZ12 18974828 ChIP-Seq MESC's Mouse | 1.89E-12 | PRC2 core |
| TCF21 26020271 ChIP-Seq SMOOTH MUSCLE Human | 1.90E-12 | TX factor - basic helix-loop-helix |
| RACK7 27058665 Chip-Seq MCF-7 Human | 1.78E-11 | TX regulator - RACK receptor - PRC2 regulator |
| SUZ12 27294783 Chip-Seq ESC's Mouse | 3.39E-11 | PRC2 core |
| EGR1 20690147 ChIP-Seq ERYTHROLEUKEMIA Human | 4.40E-11 | TX regulator - C2H2 zink finger |
| SUZ12 18692474 ChIP-Seq MEFs Mouse | 9.20E-11 | PRC2 core |
| KDM2B 26808549 Chip-Seq K562 Human | 2.20E-10 | PRC1 core |
| EZH2 27294783 Chip-Seq ESC's Mouse | 2.20E-10 | PRC2 core |
| JARID2 20075857 ChIP-Seq MESC's Mouse | 6.20E-10 | PRC2.2 facultative subunit |
| RNF2 18974828 ChIP-Seq MESC's Mouse | 6.31E-10 | PRC1 core |
| EZH2 18974828 ChIP-Seq MESC's Mouse | 6.31E-10 | PRC2 core |
| WT1 20215353 ChIP-ChIP NEPHRON PROGENITOR Mouse | 6.31E-10 | TX factor |
| RING1B 27294783 Chip-Seq ESC's Mouse | 6.75E-10 | PRC1 core |
| SUZ12 18692474 ChIP-Seq MESC's Mouse | 1.05E-09 | PRC2 core |
| SUZ12 18555785 ChIP-Seq MESC's Mouse | 4.07E-09 | PRC2 core |
| KDM2B 26808549 Chip-Seq SUP-B15 Human | 4.83E-09 | PRC1 core |
| SRY 25088423 ChIP-ChIP EMBRYONIC GONADS Mouse | 7.06E-09 | TX factor - SRY-Box |
| KLF4 26769127 Chip-Seq PDAC-Cell line Human | 3.39E-08 | TX - Kruppel |
| RNF2 27304074 Chip-Seq ESC's Mouse | 5.77E-08 | PRC1 core |
| WT1 25993318 ChIP-Seq PODOCYTE Human | 7.32E-08 | TX factor |
| JARID2 20064375 ChIP-Seq MESC's Mouse | 9.08E-08 | PRC2.2 facultative subunit |
| SOX2 20726797 ChIP-Seq SW620 Human | 1.31E-07 | TX factor - SRY-Box |
| RARG 19884340 ChIP-ChIP MEFs Mouse | 2.46E-07 | TX factor - ligand activated |
| UBF1/2 26484160 Chip-Seq HMEC-DERIVED Human | 3.52E-07 | TX factor - RNA transcription |
| RUNX2 24764292 ChIP-Seq MC3T3 Mouse | 6.32E-07 | TX factor |
| TP53 20018659 ChIP-ChIP R1E Mouse | 9.40E-07 | TX factor - P53 family |
| SMC1 22415368 ChIP-Seq MEFs Mouse | 1.09E-06 | cohesin subunit - PRC1 regulator |
| ZNF217 24962896 ChIP-Seq MCF-7 Human | 2.05E-06 | TX factor - repressor - PRC2 regulator |
| SA1 22415368 ChIP-Seq MEFs Mouse | 3.08E-06 | cohesin subunit |
| KDM2B 26808549 Chip-Seq JURKAT Human | 3.08E-06 | PRC1 core |
| RING1B 27294783 Chip-Seq NPCs Mouse | 3.08E-06 | PRC1 core |
| KLF5 25053715 ChIP-Seq YYC3 Human | 3.08E-06 | TX - Kruppel |
| ESR2 21235772 ChIP-Seq MCF-7 Human | 4.87E-06 | TX factor - hormone activated |
| KDM2B 26808549 Chip-Seq DND41 Human | 9.18E-06 | PRC1 core |
| EOMES 21245162 ChIP-Seq HESCs Human | 1.05E-05 | TX factor - T-box |
| P300 27058665 Chip-Seq ZR-75-30cells Human | 1.49E-05 | HAT |
| JUN 26020271 ChIP-Seq SMOOTH MUSCLE Human | 1.49E-05 | TX factor - proto-oncogene |
| KLF6 26769127 Chip-Seq PDAC-Cell line Human | 2.39E-05 | TX - Kruppel |
| CJUN 26792858 Chip-Seq BT549 Human | 2.39E-05 | TX factor - proto-oncogene |
| CTCF 27219007 Chip-Seq ERYTHROID Human | 2.39E-05 | TX regulator - zink finger - PRC2 regulator |
| ATF3 23680149 ChIP-Seq GBM1-GSC Human | 2.70E-05 | TX factor - CREB |
| BACH1 22875853 ChIP-PCR HELA AND SCP4 Human | 3.63E-05 | TX factor - CNC-bZip - PRC2 regulator |
| KDM2B 26808549 Chip-Seq SIL-ALL Human | 6.35E-05 | PRC1 core |
| ZFP281 27345836 Chip-Seq ESCs Mouse | 6.35E-05 | TX factor - repressor - PRC2 regulator |
| CEBPD 21427703 ChIP-Seq 3T3-L1 Mouse | 7.41E-05 | TX factor - bZIP |
| P63 26484246 Chip-Seq KERATINOCYTES Human | 0.00010 | TX factor - P53 family |
| TP63 17297297 ChIP-ChIP HaCaT Human | 0.00013 | TX factor - P53 family |
| EED 16625203 ChIP-ChIP MESC's Mouse | 0.00014 | PRC2 core |
| NFI 21473784 ChIP-Seq ESCs Mouse | 0.00016 | TX factor - CTF/NF-I family |
| EZH2 27304074 Chip-Seq ESCs Mouse | 0.00021 | PRC2 core |
| ELF3 26769127 Chip-Seq PDAC-Cell line Human | 0.00025 | TX factor |
| LXR 22292898 ChIP-Seq THP-1 Human | 0.00025 | TX factor - ligand activated |
| ESR1 21235772 ChIP-Seq MCF-7 Human | 0.00033 | TX factor - hormone activated |
| BRD4 25478319 ChIP-Seq HGPS Human | 0.00037 | TX factor - bromodomain |
| SMC3 22415368 ChIP-Seq MEFs Mouse | 0.00037 | cohesin subunit - PRC1 regulator |
| CLOCK 20551151 ChIP-Seq 293T Human | 0.00037 | TX factor - regulation of circadian rhythms |
| CTCF 27219007 Chip-Seq Bcells Human | 0.00037 | TX regulator - zinc finger - PRC2 regulator |
| SUZ12 16625203 ChIP-ChIP MESC's Mouse | 0.00057 | PRC2 core |
| CREB1 26743006 Chip-Seq LNCaP-abl Human | 0.00057 | TX factor - CREB |
| CTCF 21964334 Chip-Seq Bcells Human | 0.00057 | TX regulator - zinc finger - PRC2 regulator |
| TP53 23651856 ChIP-Seq MEFs Mouse | 0.00071 | TX factor - P53 family |
| KLF4 18358816 ChIP-ChIP MESC's Mouse | 0.00091 | TX - Kruppel |
| SOX9 24532713 ChIP-Seq HFSC Mouse | 0.00097 | TX factor - SRY-Box |
| NUCKS1 24931609 ChIP-Seq HEPATOCYTES Mouse | 0.00097 | TX regulator - dna repair |

**Supplementary Table III.** Epigenomic datasets identified with 723 upregulated transcripts in human Tau-KO cells (Adj P<.01). Analysis performed August 4^th^ 2022.

| Epigenomics Roadmap HM ChIP Term | Adj P | Comment |
| --- | --- | --- |
| H3K27me3 H1 | 6.4E-23 | H3K27me3 |
| H3K27me3 Colonic Mucosa | 1.2E-18 | H3K27me3 |
| H3K27me3 Mobilized CD34 Primary Cells | 2.9E-16 | H3K27me3 |
| H3K27me3 Fetal Brain | 1.8E-15 | H3K27me3 |
| H3K27me3 iPS-20b | 1.2E-11 | H3K27me3 |
| H3K27me3 iPS DF 19.11 | 7.3E-11 | H3K27me3 |
| H3K27me3 CD34 Primary Cells | 3.1E-10 | H3K27me3 |
| H3K27me3 H9 | 4.2E-10 | H3K27me3 |
| H3K27me3 CD8 Memory Primary Cells | 3.1E-09 | H3K27me3 |
| H3K27me3 iPS DF 6.9 | 1.4E-08 | H3K27me3 |
| H3K27me3 CD4+ CD25- CD45RA+ Naive Primary Cells | 2.2E-08 | H3K27me3 |
| H3K27me3 CD8 Naive Primary Cells | 3.2E-08 | H3K27me3 |
| H3K27me3 Rectal Smooth Muscle | 4.0E-08 | H3K27me3 |
| H3K27me3 H1 BMP4 Derived Mesendoderm Cultured Cells | 9.7E-08 | H3K27me3 |
| H2BK20ac IMR90 | 1.7E-07 | H2BK20ac |
| H3K27me3 CD4+ CD25int CD127+ Tmem Primary Cells | 2.6E-07 | H3K27me3 |
| H3K27me3 Duodenum Mucosa | 3.1E-07 | H3K27me3 |
| H3K27me3 iPS-15b | 1.3E-06 | H3K27me3 |
| H3K27me3 H1 BMP4 Derived Trophoblast Cultured Cells | 1.9E-06 | H3K27me3 |
| H3K4me1 CD4+ CD25- CD45RA+ Naive Primary Cells | 9.5E-06 | H3K4me1 |
| H3K27me3 Rectal Mucosa | 1.6E-05 | H3K27me3 |
| H3K4me1 H1 | 3.2E-05 | H3K4me1 |
| H3K4me1 Brain Germinal Matrix | 3.4E-05 | H3K4me1 |
| H2BK20ac H1 | 3.5E-05 | H2BK20ac |
| H3K27me3 Brain Germinal Matrix | 4.6E-05 | H3K27me3 |
| H3K4me1 IMR90 | 5.8E-05 | H3K4me1 |
| H3K27me3 CD4+ CD25- CD45RO+ Memory Primary Cells | 1.4E-04 | H3K27me3 |
| H3K27me3 Pancreatic Islets | 1.9E-04 | H3K27me3 |
| H3K27me3 CD3 Primary Cells | 2.4E-04 | H3K27me3 |
| H3K27me3 Neurosphere Cultured Cells Cortex Derived | 2.5E-04 | H3K27me3 |
| H2BK12ac IMR90 | 2.7E-04 | H2BK12ac |
| H3K4me1 CD4+ CD25- Th Primary Cells | 2.7E-04 | H3K4me1 |
| H3K4me1 Fetal Brain | 2.8E-04 | H3K4me1 |
| H3K27me3 CD4 Naive Primary Cells | 3.2E-04 | H3K27me3 |
| H3K27me3 Stomach Mucosa | 4.3E-04 | H3K27me3 |
| H3K27me3 CD4+ CD25+ CD127- Treg Primary Cells | 5.2E-04 | H3K27me3 |
| H3K27me3 Fetal Lung | 7.6E-04 | H3K27me3 |
| H3K27me3 CD4+ CD25- Th Primary Cells | 8.1E-04 | H3K27me3 |
| H3K27me3 Duodenum Smooth Muscle | 0.0011 | H3K27me3 |
| H2BK15ac IMR90 | 0.0018 | H2BK15ac |
| H3K27me3 Brain Hippocampus Middle | 0.0021 | H3K27me3 |
| H3K27me3 CD4 Memory Primary Cells | 0.0028 | H3K27me3 |
| H2BK120ac IMR90 | 0.0031 | H2BK120ac |
| H3K27me3 Brain Substantia Nigra | 0.0031 | H3K27me3 |
| H3K4me1 iPS DF 6.9 | 6.6E-03 | H3K4me1 |

**Supplementary Table IV:**
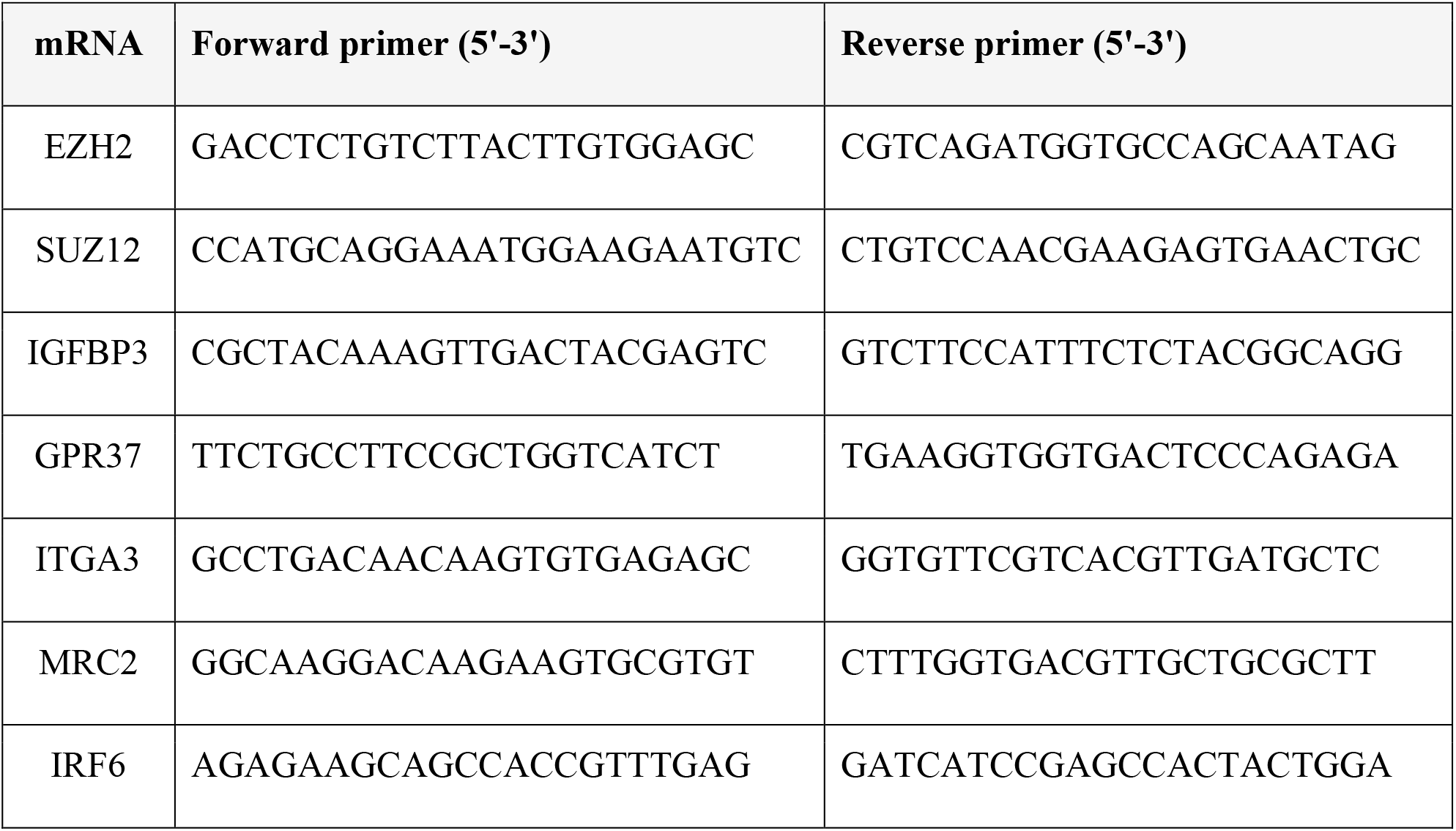
qPCR primers.

